# An online tool for calculating initial rates from continuous enzyme kinetic traces

**DOI:** 10.1101/700138

**Authors:** Michael D. Olp, Kelsey S. Kalous, Brian C. Smith

## Abstract

A computer program was developed for semi-automated calculation of initial rates from continuous kinetic traces during the evaluation of Michaelis-Menten and EC_50_/IC_50_ kinetic parameters from high-throughput enzyme assays. The tool allows users to interactively fit kinetic traces using convenient browser-based selection tools, ameliorating tedious steps involved in defining linear ranges in general purpose programs like Microsoft Excel while still maintaining the simplicity of the “ruler and pencil” method of determining initial rates. As a test case, we quickly and accurately analyzed over 500 continuous enzyme kinetic traces resulting from experimental data on the response of Sirt1 to small-molecule activators. For a given titration series, our program allows simultaneous visualization of individual initial rates and the resulting Michaelis-Menten or EC_50_/IC_50_ kinetic model fit. In addition to serving as a convenient program for practicing enzymologists, our tool is also a useful teaching aid to visually demonstrate in real-time how incorrect initial rate fits can affect calculated steady-state kinetic parameters. For the convenience of the research community, we have made our program freely available online at https://continuous-enzyme-kinetics.herokuapp.com/continuous-enzyme-kinetics.

## Background

Continuous enzyme kinetic assays allow rapid acquisition of large numbers of kinetic traces. Therefore, data analysis often becomes the bottleneck of high-throughput enzymatic screening pipelines. In cases where IC_50_/EC_50_ values or the Michaelis-Menten parameters *V* _max_ (or *k* _cat_) and *K* _M_ are of principle interest, reduction of kinetic traces to solely the initial velocities avoids error arising from assumptions involved in analyzing the entire kinetic trace [1]. The two primary methods for determining initial velocities from kinetic traces are *(i)* “ruler and pencil” estimation of the early linear portion of the curve and *(ii)* methods using integrated forms of kinetic equations [2, 3, 4]. Currently available programs such as FITSIM [5], DYNAFIT [6], ENZO [7], PCAT [8] and KinTek offer sophisticated routines for fitting entire kinetic traces curves using approaches falling under method *(ii)*. These software packages are very useful for selecting among complex enzymatic models and analyzing experiments carried out under conditions that may not satisfy the assumptions associated with Michaelis-Menten kinetics [9, 10], for example measuring catalysis inside cells. However, the complexity offered by these tools is often not required when analyzing *in vitro* experiments where sufficient data can be collected under initial rate conditions, making them inefficient for many high-throughput applications. As a result, method *(i)*remains by far the most used method for calculating initial rates from continuous kinetic traces.

Despite the simplicity of the “ruler and pencil” method, manual inspection and selection of a linear range from each individual kinetic trace using graphing programs such as Microsoft Excel and GraphPad can be pain-staking, time consuming and susceptible to subjective human error, particularly when low substrate concentrations result in significant curvature. As opposed to method *(ii)*, to our knowledge no software exists aimed at expediting the simple selection of initial rates by inspection. To expedite our own initial rate analyses, we developed a tool for semi-automated initial rate calculations that maintains the convenience of “ruler and pencil” estimation while increasing the accuracy of this method by allowing for rapid and user interactive visualization of initial rate fits. For the convenience of the research community, we have converted our program into a publicly-available browser-based tool (https://continuous-enzyme-kinetics.herokuapp.com/continuous-enzyme-kinetics) to which users can upload series of kinetic traces in comma separated value (CSV) format and download the resulting table of initial rates for further analysis and plotting. The tool presented here has several advantages over other available software packages for analyzing enzyme kinetics experiments in that it is free and open source, does not require any downloads prior to use (Table 1). In addition, our program includes a plot of the Michaelis-Menten (or IC_50_/EC_50_) fit for the uploaded experiment that is automatically updated based on user interaction with the time ranges used to calculate the initial rates (Table 1). As a result, the tool also serves as a useful teaching aid when demonstrating how incorrect fitting of an initial rate from a kinetic trace can affect the overall steady-state kinetic parameters calculated from a given experiment.

**Table 1.**
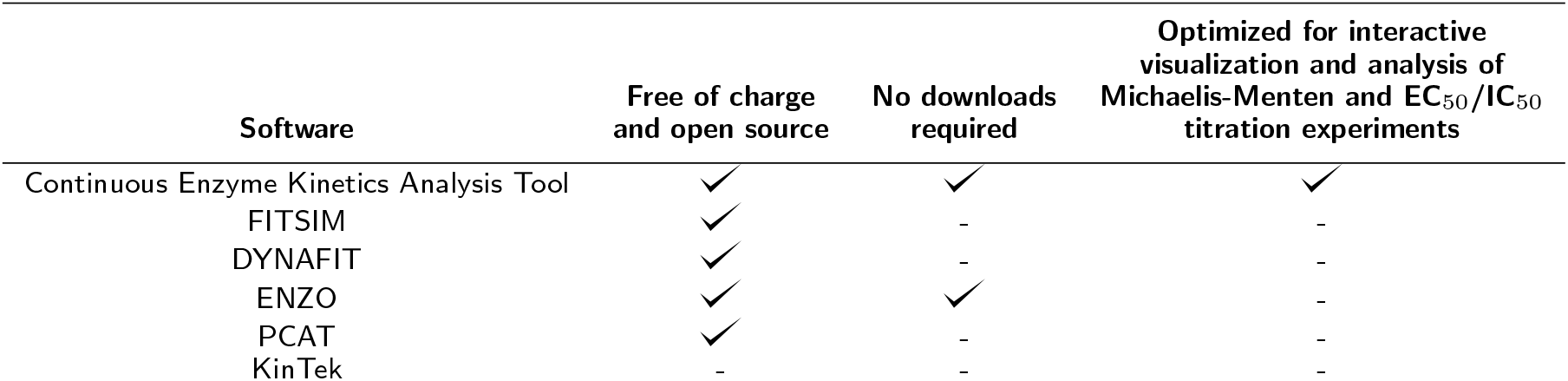
Comparison of available software.

## Implementation

### Web-based tool for continuous enzyme kinetic analysis

All calculations are carried out in Python using numpy, and both linear and non-linear regression is performed using the curve fit method from the scipy.optimize module. In linear mode, slopes corresponding to initial rates are determined using a straight line fit to a user-specified segment of the kinetic trace. When using logarithmic mode, selected kinetic traces are fit to a logarithmic approximation of the integrated Michaelis-Menten equation defined by

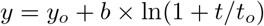

where y_o_ is the background signal, t_o_ *>* 0 is the scale of the logarithmic curve, and b *>* 0 is a shape parameter [3]. The kinetic trace slope corresponding to the initial rate is equal to the first derivative of the logarithmic fit when t = 0. Kinetic trace slopes are converted to initial rates using a user-defined transform equation entered into a text box. The source code for the online tool is available at https://github.com/SmithLabMCW/continuous-enzyme-kinetics.

## Results and Discussion

Publicly-available webtool for semi-automated and interactive initial rate calculations. Continuous enzyme kinetic traces for titration experiments are uploaded in CSV format using the green button labeled “Upload Local File” at the top of the page (Figure 1a). Each CSV file should have one column containing time in seconds or minutes. The remaining CSV columns should contain time-course data, where each column heading contains a number corresponding to titrant concentration (an example CSV file is included as Appendix B and at https://github.com/SmithLabMCW/continuous_enzyme_kinetics/blob/master/continuous-enzyme-kinetics/test.csv). Depending on the type of experiment being analyzed, users can choose to fit datasets in Michaelis-Menten, EC_50_/IC_50_ or high-throughput screening (HTS) modes using the dropdown menu labeled “Choose Model” (Figure 1b). Upon file upload, all kinetic traces are automatically fit to a straight line that maximizes slope magnitude (Figure 1b) the model fit for the dataset is plotted to the right of the selected trace (Figure 1e), and the initial rate and model fit values with propagated errors are listed in data tables (Figure 1h). Users can select individual kinetic traces using the dropdown menu “Y Axis Sample” (Figure 1b) and manually refit subsets of the time-course data to obtain random residual distributions using the range slider tool or by entering start and end times in the “Start Time” and “End Time” text boxes (Figure 1f). Upon refitting an individual kinetic trace, the model fit plot (Figure 1g) and the data tables (Figure 1h) are automatically updated. Users may subtract the slope of a blank sample from the rest of the dataset using the Select Blank Sample for Subtraction” dropdown menu (Figure 1b). Users can also transform slope values into meaningful initial rates by entering a transform equation (signal as a function of time “x”, *e.g.* “x/*(extinction coefficient × path length × enzyme concentration)*”) in the “Enter Transform Equation” textbox (Figure 1c). Finally, when the user is satisfied with the analysis, the initial velocities listed in the table at the right can be copied to the clipboard by clicking the blue button labeled “Copy Table to Clipboard” or downloaded as a CSV file using the blue button labeled “Download Table to CSV” (Figure 1h).

**Figure 1.**
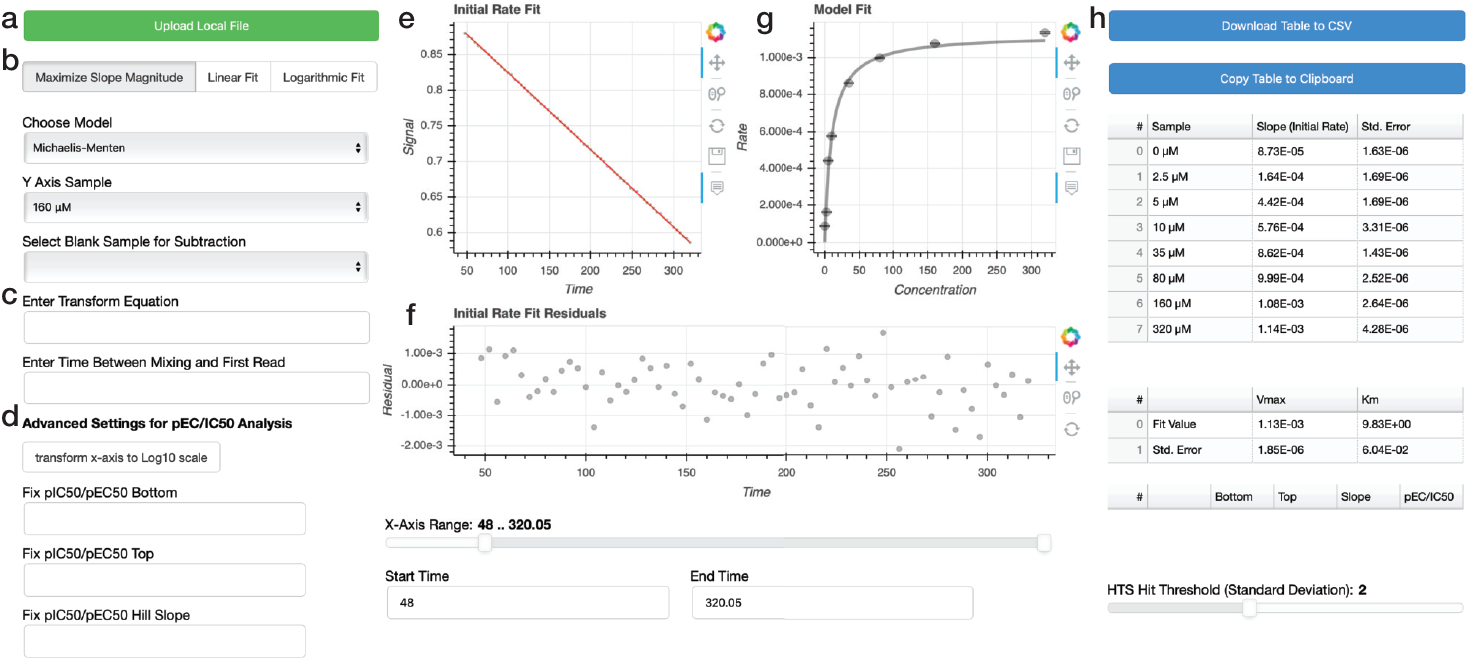
Online tool for interactive continuous enzyme kinetic trace analysis. (**a**) Click the “Upload Local File” to begin analysis of user CSV formatted data. (**b**) Click buttons to select routine for fitting the kinetic traces. The default is to maximize the slope magnitude. Use dropdown menus to select between Michaelis-Menten, IC_50_/EC_50_ and high-throughput screening (HTS) modes, choose x- and y-axis samples, and select a blank sample for subtraction. (**c)** Use the text boxes to transform kinetic trace slope values to initial rates when using linear fitting mode, as well as enter a time delay between mixing and first measurement (used in logarithmic fitting mode only).(**d**) Advanced settings for transforming the input x-axis values from a linear to a log scale for analysis and plotting, fixing the bottom and/or top of the fitted curve to a particular value, and fixing the Hill slope of the fitted curve to a particularly value (typically 1). (**e**) Representative continuous enzyme kinetic trace (grey) with initial rate fit (red) corresponding to the selected y-axis sample. (**f**) Plot of the residuals from the kinetic trace initial rate fit in (**e**). The slider and text enter boxes both allow the user to optimize the time domain of the fit to obtain a random residual distribution. (**g**) Plot of a Michaelis-Menten model fit of initial rates. (**h 7**) Data table containing initial rate values and model fit values with propagated errors. Use the “Download Table to CSV” or “Copy Table to Clipboard” buttons to export initial rate values from the data table.

### Kinetic trace fitting routines

Two of the most commonly used approaches for determining initial velocities from continuous enzyme kinetic traces include *(i)* “ruler and pencil” estimation of the early linear portion of the curve and *(ii)* methods using integrated forms of kinetic equations [2, 3, 4]. The most mathematically rigorous methods for calculating accurate initial velocities fall under method *(ii)*. These integrated kinetic equations are particularly important to consider when the portion of the kinetic trace corresponding to the initial rate is difficult to measure, as in situations where substrate concentrations are below the *K* _M_ value of an enzyme. Early methods using the integrated Michaelis-Menten equation are highly sensitive to assumptions regarding reaction reversibility, product inhibition, and enzyme inactivation and stability [1]. More recent methods treating initial and final theoretical substrate concentrations as parameters in non-linear regression [2, 3, 4] eliminate these assumptions from the fitting process and have greatly increased the applicability of the integrated Michaelis-Menten equation for calculating initial velocities from kinetic traces. Nevertheless, the simplicity of estimating a straight-line has made method *(i)*by far the most prevalent means of calculating initial rates from kinetic traces.

Our program first generates an initial rate prediction using linear regression to maximize the first derivative of the kinetic traces smoothed by cubic spline interpolation. Using this method, linear initial rate estimations are automatically generated for an entire enzyme/substrate titration (Figure 2a-h) and assessed for adherence to the Michaelis-Menten or EC_50_/IC_50_equations as defined in the “Choose Model” dropdown menu (Figure 1b). To avoid error arising from erroneous fitting of kinetic artifacts, the tool allows users to interactively re-assign the time range used for fitting using the range slider tool or by entering start and end times in the“Start Time” and “End Time” text boxes (Figure 1f). Upon manual selection of a new time range for the selected kinetic trace by the user, a new initial rate is calculated, and this change is automatically reflected in the overall kinetic model fit (Figure 1g) and data tables (Figure 1h). During the refitting process, the user can select from three fitting modes by clicking on the buttons labeled “Maximize Slope Magnitude”, “Linear Fit” and “Logarithmic Fit” (Figure 1b). “Maximize Slope Magnitude” mode is the default as it is used in the automatic initial rate estimation described above. “Linear Fit” mode is equivalent to “ruler and pencil” initial rate estimation where a straight line is fit to the user-selected portion of the kinetic trace. “Logarithmic Fit” mode is an implementation of the logarithmic approximation of the integrated Michaelis-Menten equation described recently by Lu and coworkers[3]. This method is particularly useful to avoid under-estimation of initial rates from kinetic traces where an initial linear segment cannot be satisfactorily identified. If there is a significant time delay between initiating the enzyme-catalyzed reaction and the first sample read, a time value can be entered into the text box labeled “Enter Time Between Mixing and First Read” (Figure 1c) to extrapolate initial rate calculations back to the exact time of mixing (Note, the value entered in this text box is used in the calculation only when logarithmic fitting mode is selected). Throughout the fitting process, users should strive to obtain a random distribution of points in the kinetic trace fit residual plot located directly below the kinetic trace (Figure 1f). In addition, users can dynamically assess how changes in initial rate calculations for each kinetic trace affect the overall fit of a titration to the Michaelis-Menten or EC_50_/IC_50_equations.

**Figure 2.**
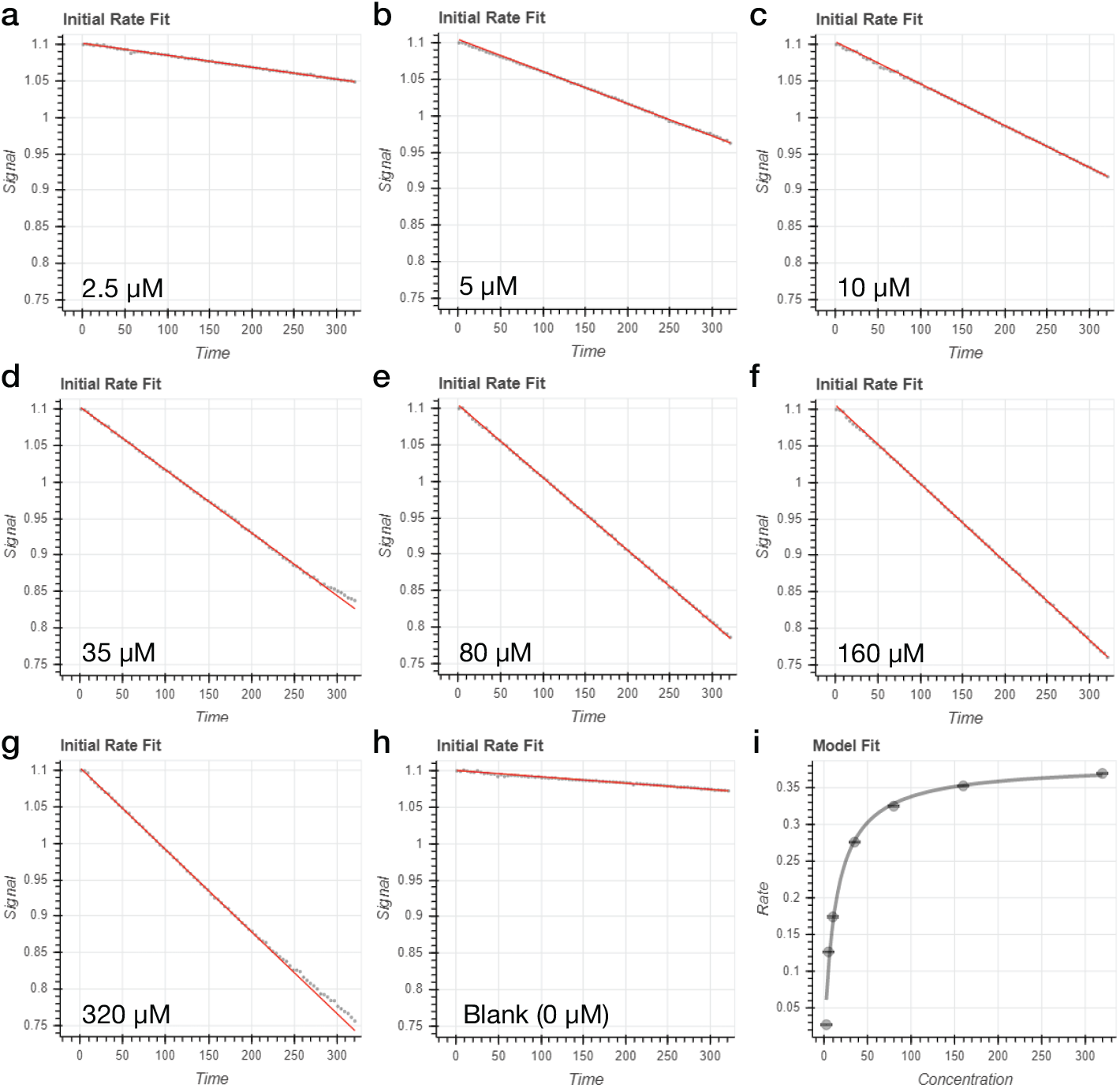
Automated determination of steady-state kinetic parameters. (**a-h**) Automated fits (red lines) generated by the interactive tool for continuous enzyme kinetic traces from a representative dataset using substrate concentrations ranging from 0 to 320 *µ*M (grey points). (**i**) Michaelis-Menten plot generated by the interactive tool for continuous enzyme kinetics.

### EC_50_/IC_50_ and HTS fitting modes

In addition to Michaelis-Menten kinetics, our software is also optimized to perform analysis of datasets resulting from EC_50_/IC_50_ (Figure 3a/b) and HTS (Figure 3c/d) kinetic experiments. In each case, initial rates are determined in the same manner as described above for Michaelis-Menten kinetics. When working in EC_50_/IC_50_ mode, changes in initial rate values and associated errors are automatically reflected in the fit to the 4-parameter logistic model:

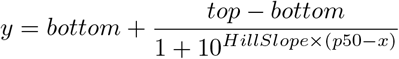

(Figure 3b). Advanced EC_50_/IC_50_ analysis settings allow users to inter-convert the x-axis between linear and Log_10_ scale as well as fix the top, bottom and Hill Slope regression values (Figure 3b). The data table containing the four regression parameters and propagated errors is automatically updated throughout interactive initial rate fitting (Figure 3b).

**Figure 3.**
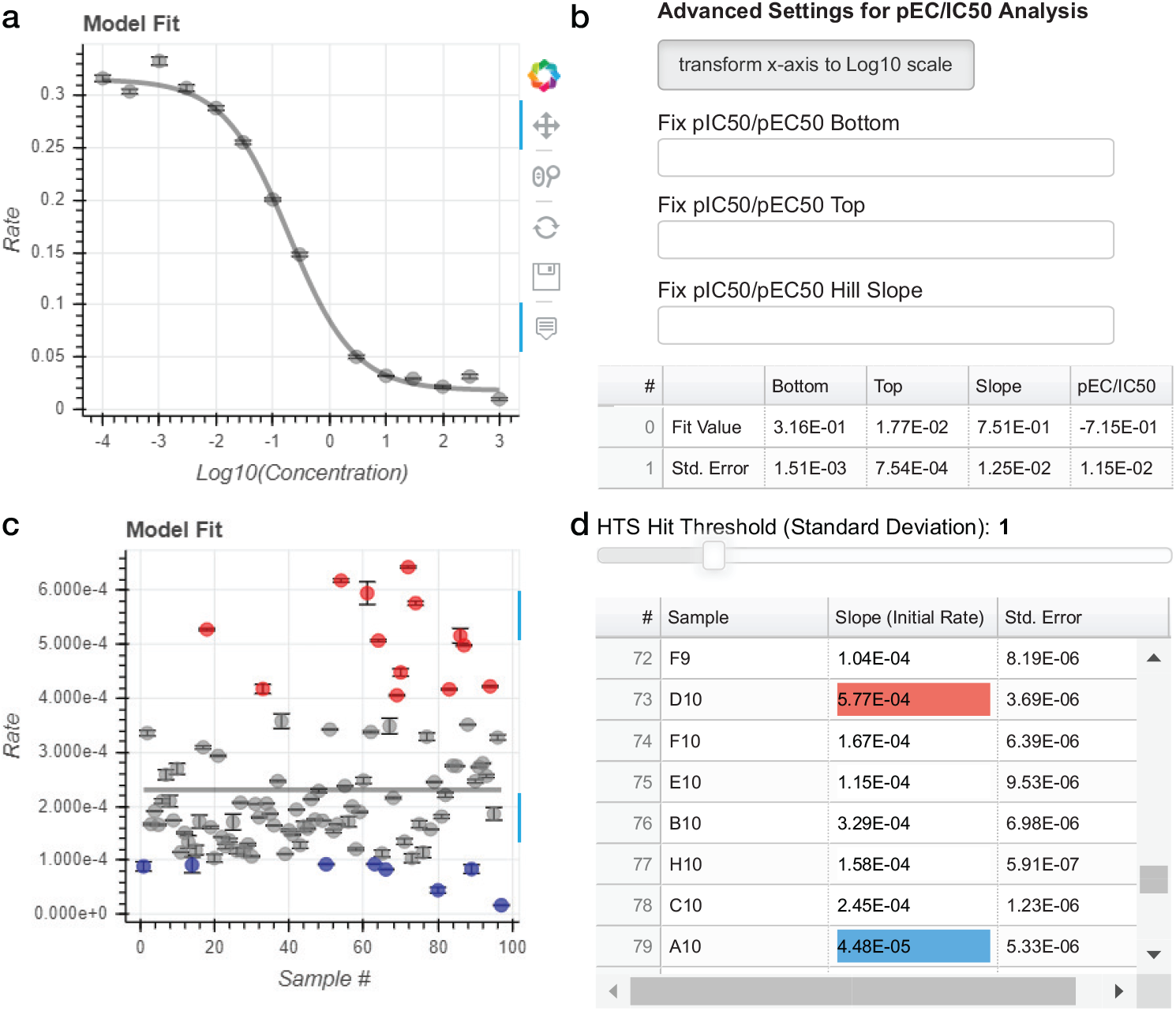
EC_50_/IC_50_ and HTS analysis modes. (**a**)Plot of a representative IC_50_ model fit of initial rates. (**b**) Widgets for choosing advanced EC_50_/IC_50_ analysis settings allow users to convert the x-axis to Log_10_ scale and fix regression parameters. The data table displays fit values and propagated errors for the 4-parameter logistic model. (**c**) Plot displaying HTS analysis of initial rates. (**d**) The data table displays initial rates and associated errors for all samples uploaded and highlights cells corresponding to samples with initial rates above (red) or below (blue) the standard deviation threshold defined by the slider (here set to 1 standard deviation from the mean initial rate).

Also implemented is an HTS mode, within which an unlimited number of samples (e.g. activator/inhibitor screening in 96-or 384-well plate format) can be uploaded and fit to determine initial rates. When analyzing data in HTS mode, a straight horizontal line is plotted to represent the mean initial rate of the data set and samples associated with initial rates either above (red) or below (blue) a user-defined standard deviation threshold from the mean are highlighted on the model fit plot (Figure 3c) as well as in the data table (Figure 3d).

### Case-study: Interactive dataset fitting as a visual teaching aid

A key utility of this tool is to teach students or train new laboratory members to fit continuous enzyme kinetic data. In particular, the tool can be used to interactively demonstrate proper identification and selection of the initial rate component of a kinetic trace, as well as the consequences of incorrect identification of initial rates (Figure 4). Fitting a line segment temporally downstream of the initial rate segment results in underestimation of rate (Figure 3a-c). In Michaelis-Menten analysis, underestimation of rates, especially from kinetic traces where the concentration of substrate is low, results in underestimation of the *K* _M_ value for the enzyme (Figure 3d/e). This phenomenon can be demonstrated by manually using the sliders to intentionally select an incorrect line segment from continuous kinetic data (Figure 1f). As the software automatically updates the overall fit of the entire data set (*K* _M_ and *v_max_/k_cat_* values) as adjustments are made, students and trainees are able to immediately visualize the impact of underestimating an initial rate on the overall fit of a Michaelis-Menten curve (Figure 3d/e). Adjustment of the sliders to fit different components of a curve, and rapid integration of the adjusted rates into the overall fit, allows for fluid demonstration of initial rate fitting in the context of a lecture, which otherwise would be discontinuous and cumbersome using programs such as GraphPad or Microsoft Excel.

**Figure 4.**
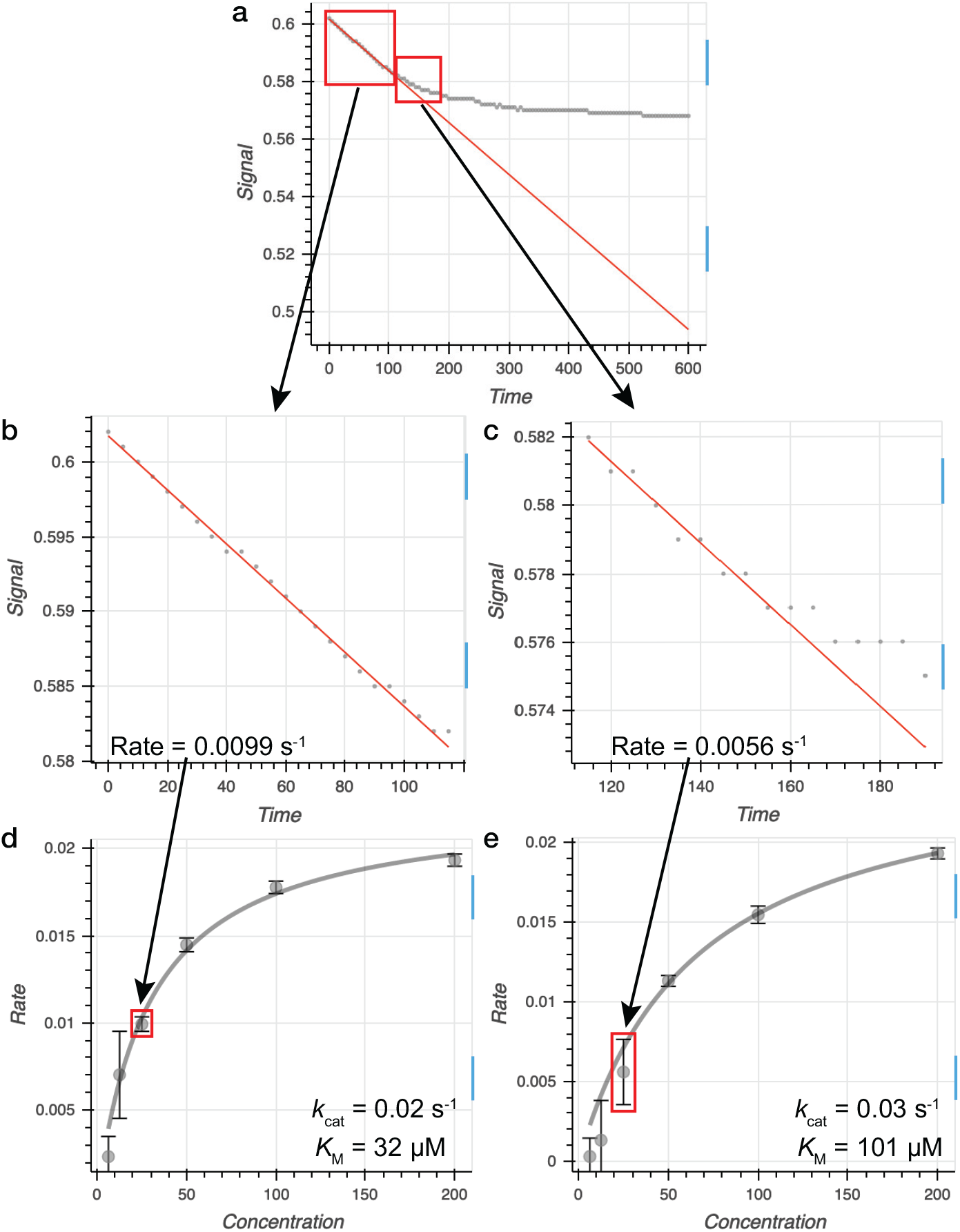
Interactive dataset fitting as a visual teaching aid. (a) A representative continuous enzyme kinetic trace where either (**b**) the initial linear rate is fit appropriately yielding (**d**) accurate initial rates for the Michaelis-Menten fit or (**c**) the kinetic trace is fit after the initial rate has passed typically yielding (**e**) a Michaelis-Menten fit with an inaccurate *K* _M_ value higher than the actual *K* _M_value.

### Case-study: Rapid determination of initial rates and steady-state kinetic parameters for Sirt1 mutants with small-molecule activators

The Sirt1 deacetylase [11] protects against aging-related disorders [12, 13, 14], and Sirt1 activators (STACs) [15, 16, 17, 18, 19] are sought as therapeutics. Resveratrol and other STACs (Figure S1) activate Sirt1 by binding the Sirt1 *N*-terminal domain (residues 183-230) [17] and lower the *K* _M_ value of a subset of acetylated substrates [15, 16, 18]. Crystallization and mutagenesis studies suggest that residues in the Sirt1 catalytic core (residues 244-498) [17] may also mediate conformational changes critical for Sirt1 activation [15, 17, 18], but the importance of *N*-terminal domain versus catalytic core residues in Sirt1 activation had not been investigated.

To test the ability of our online tool to rapidly determine steady-state kinetic parameters from continuous enzyme kinetic traces, six Sirt1 mutations (I223A, I223R, E230K, D292A, F414A, and R446E) were generated based on previous crystallographic, hydrogen-deuterium exchange, and kinetic studies [15, 17, 18]. The activity of each mutant was screened in the presence or absence of resveratrol or STAC1 (Figure S1) using a high-throughput continuous enzyme-coupled assay for sirtuins [20]. Given the combinatorial nature of this study (seven different Sirt1 constructs, seven substrate concentrations, two Sirt1-activating compounds, and a minimum of three experimental replicates), a large volume of kinetic data was generated, providing an excellent test case of our online tool for semi-automated processing of steady-state kinetic data. This tool was used to quickly and accurately process over 500 kinetic traces obtained from Michaelis-Menten titrations of an acetylated p53-based peptide [15, 17, 18]. Kinetic parameters (*k* _cat_, *K* _M_, and *k* _cat_/*K* _M_) were assessed and compared to determine the relative impact of each mutation on Sirt1 activation. Our data indicate that I223, D292, F414 and R446 are required for both resveratrol- and STAC1-mediated Sirt1 activation. Interestingly, the E230K mutant was selectively activated by STAC1, indicating the Sirt1 binding site and/or activation mechanism is not identical for resveratrol and STAC1. Supplemental discussion is included in Appendix A.

## Conclusions

To increase the speed and accuracy of the data analysis stage of continuous enzyme kinetic assays, a publicly available web-based tool was developed for semi-automated and interactive continuous enzyme kinetic trace analysis. Our program offers several advantages over other available software packages for analyzing continuous enzyme kinetics experiments in that it is free, web-based and optimized interactive and intuitive analysis of Michaelis-Menten, EC_50_/IC_50_ and HTS data sets. As a case study for this tool, a comprehensive kinetic screen using a continuous enzyme-coupled assay for sirtuins [20] was conducted to examine the relative contributions of five Sirt1*N*-terminal andcatalytic domain residues to resveratrol and STAC1-induced enhancement of Sirt1 catalytic efficiency. In addition to helping researchers increase the efficiency of kinetic trace analyses, the interface serves as a useful teaching tool for demonstrating the link between accurate initial rate determination and calculation of Michaelis-Menten and EC_50_/IC_50_kinetic parameters.

## Availability and requirements

**Project name:**

Continuous Enzyme Kinetics Analysis Tool

**Project home page:**

https://continuous-enzyme-kinetics.herokuapp.com/continuous-enzyme-kinetics

**Archived version:**

N/A

**0.1 Operating system(s):**

Platform independent

**Programming language:**

Python, Java

**License:**

N/A

**Any restrictions to use by non-academics:**

N/A

### Declarations

Ethics approval and consent to participate Not applicable

### Consent for publication

Not applicable

### Availability of data and material

The program described here is freely available at https://continuous-enzyme-kinetics.herokuapp.com/continuous-enzyme-kinetics. All source code is present in the associated GitHub repository located at https://github.com/SmithLabMCW/continuous_enzyme_kinetics. While no uploaded data is saved by this application, users concerned about privacy can download the associated GitHub repository (https://github.com/SmithLabMCW/continuous_enzyme_kinetics.git) and run the application locally (see https://github.com/SmithLabMCW/continuous_enzyme_kinetics for usage instructions).

### Competing interests

Not applicable

## Funding

This work was supported by the National Institutes of Health (R35GM128840 to B.C.S. and F31DK117588 to K.S.K.), the National Science Foundation (CHE-1708829 to B.C.S.), the American Cancer Society (14-247-29-IRG and 86-004-26-IRG to B.C.S.), the American Diabetes Association (1-18-IBS-068 to B.C.S.), and the American Heart Association (15SDG25830057 to B.C.S.).

## Authors’ contributions

### Acknowledgements

We thank members of the Smith laboratory for their input during the development of this software.

## Additional Files

Appendix A — Supplemental materials, methods, discussion, and references.

Table S1 (Sirt1 mutagenesis primers), Figure S1 (Sirt1 mutant *k*_cat_ and *K* _M_ values varying acetylated peptide in the presence of resveratrol and STAC1).

Appendix B — Sample continuous kinetic trace input data file.

