## Appendix A for "An online tool for calculating initial rates from continuous enzyme kinetic traces"

Michael D. Olp[a], Kelsey S. Kalous[a], Brian C. Smith[a*]
[a]Department of Biochemistry, Medical College of Wisconsin

Appendix A

**Supplemental Materials & Methods**

***Materials.*** Resveratrol, (L)-glutamic dehydrogenase (from bovine liver), $NAD^+$, and trifluoroacetic acid were purchased from Sigma-Aldrich (Milwaukee, WI). Fmoc amino acids, α-ketoglutaric acid, and NADH were purchased from Chem-Impex (Wood Dale, IL). Rink-amide 4-methylbenzhydrylamine resin was purchased from Novabiochem. Ni-NTA superflow resin was purchased from 5 PRIME (Hilden, Germany). STAC1 was generous gift from GlaxoSmithKline.

***Solid-phase peptide synthesis.*** A 5-mer acetyl-lysine peptide based on p53 (p53Wac: $H_2N$-RHKK(acetyl)W-$CONH_2$) was synthesized using tBu/Fmoc solid-phase peptide synthesis on an Applied Biosystems ABI 433A peptide synthesizer (FastMoc 0.1 mmol). Fmoc amino acids were coupled to 100-200-mesh Rink-amide 4-methylbenzhydrylamine resin. Protecting groups utilized were: boc for lysine and tryptophan, trityl for histidine, and 2,2,4,6,7-pentamethyldihydrobenzofuran-5-sulfonyl for arginine. Amino acids were coupled using 10 equiv of activated amino acid, 9 equiv of 2-(1H-benzotriazol-1-yl)-1,1,3,3-tetramethylluronium hexafluorophosphate, 9 equiv of hydroxybenzotriazole, and 20 equiv of *N,N*-diisopropylethylamine. Following completion of the synthesis, resin was rinsed with dichloromethane and dried. Peptide was then deprotected and cleaved from the resin using 95% (v/v) TFA, 2.5% (v/v) $H_2O$, and 2.5% (v/v) triisopropylsilane. Peptide was recovered by precipitation with cold (-20 °C) diethyl ether and centrifugation at 4,200 × g. Pelleted peptide was washed twice by resuspension in cold diethyl ether and subsequent centrifugation. Precipitated peptide was dried, redissolved in $H_2O$, frozen in liquid $N_2$, and lyophilized. Crude peptide was further purified via semipreparative reversed phase HPLC on a μBondapak C18 column (Waters, 3.9 × 300 mm) using an Agilent 1100 series HPLC. Peptide was eluted using a gradient of 0-80% (v/v) acetonitrile in water with 0.1% (v/v) TFA. Collected fractions were frozen in liquid $N_2$, lyophilized, and the masses of final products confirmed via direct injection electrospray ionization-mass spectrometry (QExactive, Thermo Scientific). The observed mass matched the predicted mass for the p53Wac peptide (calculated for $C_{39}H_{61}N_{14}O_7$ $[M+H]^+$: 837.48422, found: 837.48619). Peptide stocks were made in double deionized water and concentrations determined spectrophotometrically using the extinction coefficient for tryptophan (5.6 $mM^{-1}cm^{-1}$).

***Site-directed mutagenesis.*** Plasmids coding for Sirt1 I223A, I223R, E230K, D292A, F414A, and R446E were generated via site-directed mutagenesis of a pET28a-LIC plasmid coding for wild type Sirt1, amino acids 156-664. Mutagenesis primers were purchased from Integrated DNA Technologies (Coralville, IA) (Table A.1). Mutants were confirmed by DNA sequencing (Retrogen, San Diego, CA).

***Expression and purification of Sirt1.*** Sirt1 WT (156-664) (pET28a-LIC) and Sirt1 mutant constructs (I223A, I223R, E230K, D292A, F414A, and R446E) (pET28a-LIC) were expressed and purified from BL21(DE3) *E. coli* via nickel affinity chromatography as previously described[1]. Briefly, cells were transformed and grown at 37 °C in 2XYT media supplemented with 50 mg/L kanamycin to an optical density of 0.7 at 600 nm. Protein expression was induced with 0.5 mM IPTG at 16 °C overnight. Cells were harvested via centrifugation at 5,000 × g and bacterial pellets frozen at -80 °C. Pellets were thawed on ice and resuspended in 20 mM Tris buffer pH 8.0 containing 500 mM NaCl, 10% (v/v) glycerol, 5 mM β-mercaptoethanol, and 2.5 mM imidazole. Cells were lysed via sonication and insoluble debris cleared via centrifugation at 30,000 × g. Ni-NTA resin was added to cleared lysate (0.75 mL Ni-NTA resin/L culture) and rocked at 4 °C for 1 h. Bound resin was pelleted via centrifugation at 4,200 × g, resuspended in 10× resin volume of lysis buffer, and applied to a column. Resin was washed with 10× resin volume of 20 mM Tris buffer pH 8 containing 500 mM NaCl, 10% (v/v) glycerol, 5 mM β-mercaptoethanol, and 25 mM imidazole. Purified Sirt1 was eluted with 5× resin volume of 20 mM Tris buffer pH 8 containing 500 mM NaCl, 10% (v/v) glycerol, 5 mM β-mercaptoethanol, and 300 mM imidazole. Sirt1 was further purified via size-exclusion chromatography using an ENrich SEC 650 10 × 300 mm column, eluting into 10 mM HEPES buffer pH 7.5 containing 150 mM NaCl, 10% (v/v) glycerol, and 1 mM DTT. Purified Sirt1 was concentrated, aliquoted, and stored at -80 °C.

***Expression and purification of nicotinamidase.*** Maltose binding protein-tagged nicotinamidase (MBP-PncA) (pTEV6) was expressed and purified from BL21(DE3) *E. coli* via nickel affinity chromatography as previously described[2]. Briefly, cells were transformed and grown at 37 °C in 2XYT media supplemented with 50 mg/L ampicillin to an optical density of 0.7 at 600 nm. Protein expression was induced with 0.5 mM IPTG at 25 °C overnight. Cells were harvested via centrifugation at 5,000 × g and bacterial pellets frozen at -80 °C. Pellets were thawed on ice and resuspended in 20 mM potassium phosphate buffer pH 7.5 containing 500 mM NaCl and 5 mM imidazole. Cells were lysed via sonication and insoluble debris cleared via centrifugation at 30,000 × g. Ni-NTA resin was added to cleared lysate (0.75 mL Ni-NTA resin/L culture) and rocked at 4 °C for 1 h. Bound resin was pelleted via centrifugation at 4,200 × g, resuspended in 10× resin volume of lysis buffer, and applied to a column. Resin was washed with 10× resin volume of 20 mM potassium phosphate buffer pH 7.5 containing 500 mM NaCl and 25 mM imidazole. Purified MBP-PncA was eluted with 5× resin volume of 20 mM potassium phosphate buffer pH 7.5 containing 500 mM NaCl, and 300 mM imidazole. MBP-PncA was further purified via size-exclusion chromatography using an ENrich SEC 650 10 × 300 mm column, eluting into buffer containing 50 mM potassium phosphate, pH 7.5, 100 mM NaCl, and 10% (v/v) glycerol. Purified MBP-PncA was concentrated, aliquoted, and stored at -80 °C.

***Sirtuin enzyme coupled assay.*** Activity of Sirt1 was monitored under initial rate conditions using a continuous microplate assay for sirtuins and nicotinamide-producing enzymes[2]. The assay was performed at 25 °C in a reaction mixture containing 20 mM potassium phosphate pH 7.5, 2 mM $NAD^+$, 3.3 mM α-ketoglutarate, 200 µM NADH, 2 µM MBP-PncA, 2.5 units of (L)-glutamic dehydrogenase, 0.5 or 1 µM Sirt1 WT or mutants, and 6.25-200 µM p53Wac peptide. Rates were continuously monitored for 10 min at 340 nm in a 96-well clear flat bottom plate (Greiner Bio-One) using a BioTek Synergy Mx microplate reader (Winooski, VT). To assess Sirt1 activation by resveratrol and STAC1, Michaelis-Menten titrations of p53Wac peptide were performed under saturating concentrations of $NAD^+$ (2 mM), in the presence and absence of saturating concentrations of resveratrol (50 or 100 µM) or 50 µM STAC1. Reactions were initiated via addition of $NAD^+$. Data were analyzed and fit using the interactive fitting tool for continuous enzyme kinetics.

**Table S1.** Sirt1 mutagenesis primers

| Construct | Forward primer | Reverse primer |
|---|---|---|
| D292A | 5' CCCAGATCTTCCAGCTCCTCAAGCGATGTTTG 3' | 5' CAAACATCGCTTGAGGAGCTGGAAGATCTGGG 3' |
| E230K | 5' GGCAGATTGTTATTAATATCCTTTCAAAACCACC 3' | 5' GGTGGTTTTGAAAGGATATTAATAACAATCTGCC 3' |
| F414A | 5' CCAGAGATTGTGTTTGCTGGTGAAAATTTACCAGAAC 3' | 5' GTTCTGGTAAATTTTCACCAGCAAACACAATCTCTGG 3' |
| I223A | 5' GATGATATGACACTGTGGCAGGCTGTTATTAATATCC 3' | 5' GGATATTAATAACAGCCTGCCACAGTGTCATATCATC 3' |
| I223R | 5' GATGATATGACACTGTGGCAGCGTGTTATTAATATCC 3' | 5' GGATATTAATAACACGCTGCCACAGTGTCATATCATC 3' |

| | | |
|---|---|---|
| R446E | 5' GGGTCTTCCCTCAAAGTAGAACCAGTAGCACTAATTCC 3' | 5' GGAATTAGTGCTACTGGTTCTACTTTGAGGGAAGACCC 3' |

**Supplemental Discussion**

Consistent with previously-published data, wild type Sirt1 was activated by resveratrol and STAC1 (Figure S1a). Although not reaching statistical significance, p53Wac peptide $K_M$ (14 ± 5 μM) was reduced approximately 3.5-fold by resveratrol (4 ± 1 μM) and approximately two-fold (7 ± 3 μM) by STAC1 (Figure S1d) without perturbation of $k_{cat}$ (Figure S1c). The overall catalytic efficiency ($k_{cat}/K_M$) of Sirt1 WT (8,662 ± 2,264 $M^{-1}s^{-1}$) was significantly enhanced in the presence of resveratrol (23,273 ± 3,728 $M^{-1}s^{-1}$) and STAC1 (15,872 ± 3,315 $M^{-1}s^{-1}$) (Figure S1b). $k_{cat}$ values for Sirt1 mutant constructs were not enhanced (Figure S1c). p53Wac peptide $K_M$ values for Sirt1 mutant constructs were not significantly reduced, with the exception of Sirt1 D292A, which displayed significant reduction in $K_M$ by resveratrol (23 ± 9 μM), relative to the untreated enzyme (62 ± 9 μM) (Figure S1d). Examination of the overall catalytic efficiency of each construct in the presence and absence of resveratrol and STAC1 revealed robust activation of wild type Sirt1, but impaired activation of all mutant constructs, with the exception of E230K, which was significantly activated by STAC1 (20,684 ± 3,894 $M^{-1}s^{-1}$) relative to the untreated enzyme (5,514 ± 1,612 $M^{-1}s^{-1}$) (Figure S1b). These data are consistent with a critical role for each tested residue in mediating Sirt1 activation, either via formation of critical intramolecular, Sirt1-STAC, or Sirt1-substrate contacts.

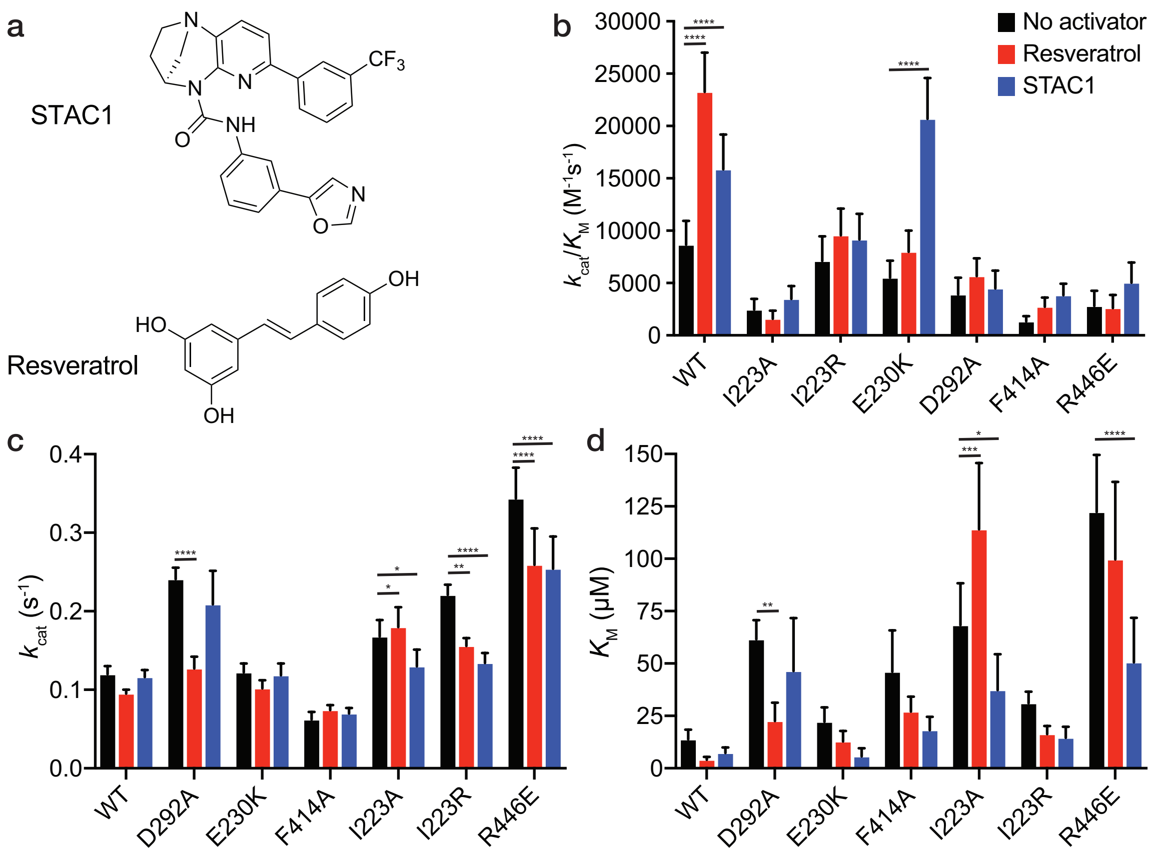


**Figure S1. Sirt1 mutant $k_{cat}$ and $K_M$ values varying acetylated peptide in the presence of resveratrol and STAC1. (a)** Chemical structures of the Sirt1 activating compounds used in this study: resveratrol and STAC1. **(b-d)** p53Wac peptide titrations were performed for Sirt1 WT and Sirt1 mutants (D292A, E230K, F414A, I223A, I223R, R446E) (0.5 or 1 μM) under saturating concentrations of $NAD^+$ (2 mM). Initial rates were fit using the interactive tool for continuous enzyme kinetics and plotted versus substrate concentration. The catalytic efficiency ($k_{cat}/K_M$) ($M^{-1}s^{-1}$), **(c)** turnover number ($k_{cat}$), and **(d)** the Michaelis constant ($K_M$) were calculated for each mutant in the presence and absence of resveratrol and STAC1. Differences in kinetic parameters were compared via two-way ANOVA (n ≥ 3) (* = $p < 0.05$, ** = $p < 0.001$, *** = $p < 0.0005$, **** = $p < 0.0001$).
